# bíogo: a simple high-performance bioinformatics toolkit for the Go language

**DOI:** 10.1101/005033

**Authors:** R. Daniel Kortschak, David L. Adelson

## Abstract

bíogo is a framework designed to ease development and maintenance of computationally intensive bioinformatics applications. The library is written in the Go programming language, a garbage-collected, strictly typed compiled language with built in support for concurrent processing, and performance comparable to C and Java. It provides a variety of data types and utility functions to facilitate manipulation and analysis of large scale genomic and other biological data. bíogo uses a concise and expressive syntax, lowering the barriers to entry for researchers needing to process large data sets with custom analyses while retaining computational safety and ease of code review. We believe bíogo provides an excellent environment for training and research in computational biology because of its combination of strict typing, simple and expressive syntax, and high performance.

## 1 Introduction

The emergence of bioinformatics as an important field of biological research has seen the proliferation of bioinformatics development toolkits (Rice *et al.*, 2000; Stajich *et al.*, 2002; Gentleman *et al.*, 2004; Knight *et al.*, 2007; Döring *et al.*, 2008; Holland *et al.*, 2008; Cock *et al.*, 2009; Goto *et al.*, 2010). Many of these toolkits are used primarily as programmatic ‘glue’, joining together functional units that are usually written in low-level high-performance languages, although many make use of high performance languages or environments internally; for examples, Pyrex/Cython in PyCogent, C in BioPerl, Biopython and Bioconductor - which also incorporates fortran code-and the Java Virtual Machine in BioRuby. This set of toolkits also includes examples written in high performance languages that have been used for building commonly used tools, for example SeqAn is used in the popular short read aligner, bowtie (Langmead *et al.*, 2009).

The use of low level languages has become necessary due to the increasingly high volumes of data encountered in modern bioinformatic analyses. However, the technical expertise required to use languages such as C, C++ and Java put their use out of the reach of many biologists, leaving Perl, Python and Ruby as the accessible options for these researchers when they wish to produce custom analytical software. Many research problems can be adequately addressed using these languages, particularly given the existence of high-performance extensions such as SciPy (http://www.scipy.org/). However, there still exist research problems where processing cannot be efficiently handled by the native language — the PyCogent authors deal with this issue by using CPU profiling techniques to identify CPU-intensive loops and rewrite ‘hot’ code in Cython/Pyrex, a Python-like language that compiles to C (Knight *et al.*, 2007; Behnel *et al.*, 2011). While this approach is effective, it requires that the authors understand two quite different languages; for although Cython is syntactically similar to Python it is, unlike Python, a statically typed language.

In addition to the issues raised above, a recent paper has highlighted the importance of good software engineering practices in scientific computing and the potential problems that may arise when these are not adopted (Wilson *et al.*, 2014). Two key principles highlighted by Wilson *et al.* are that programs should be written for people rather than machines, and that programs should be designed to account for system failure in a safe way. The first of these is at odds with the design of high performance languages such as C and C++ which expose a great deal of machine detail at the cost of readability, while the second is at odds with the design of many of the higher level, dynamically typed, languages which may allow subtle errors to silently propagate through program execution, resulting in undetected erroneous output.

## 2 New Approaches

We have chosen to approach the above problems of computational performance, ease of software development and maintenance, and computational safety by adopting a language that focuses specifically on these areas. As many of the features of blogo are a result of the design of the Go language and its tool chain we will describe the language features briefly here.

The Go language is a general purpose programming language that has been developed at Google Inc. (http://golang.org/) and as a community-based open source project. It is a compiled, strongly typed, garbage-collected language with explicit support for concurrent programming (http://golang.org/ref/spec). At approximately 50 pages in length, the language specification is short by comparison with many other languages, to allow programmers to more easily hold a complete understanding of the language’s defined behavior. The language is based on design principles intended to promote rapid and efficient software development and reduce the barriers to entry for new developers — language features should be ‘simple’, ‘orthogonal’, ‘succinct’ and ‘safe’ (R. Pike, JAOO Conference 2010). These characteristics, the language implementation and its supporting tools are well aligned with the principles for scientific computing described by Wilson *et al.* (2014): excellent support for documentation and testing; rapid write-compile-debug cycles; language-level support for concurrent processing and interprocess communication; tools for performance profiling and data race detection in concurrently executed code; and compile-time type safety in conjunction with ‘duck’ typing through interface types.

Go code is formatted mechanically to a consistent form using a tool included in the tool chain; this form, and other aspects of the Go grammar and coding style promote short lines, favoring code readability (Buse and Weimer, 2010), an important aspect in code maintainability and peer review.

Go source code and documentation are intimately tied. The Go tool chain includes an automated documentation generator that produces textual and hyper-textual documentation from source code commentary and function signatures. In the hypertext form, documentation is linked directly to source definitions via navigable hyper-links. These features provide strong connections between source and documentation and so aid development. In addition to this, the language tool chain supports inclusion of example code in the documentation. These examples are executed when running test suites, ensuring documentation parallels implementation. There is a strong culture for inclusion of good documentation and example code within the Go community, a practice that eases adoption of the language and its libraries.

As well as providing excellent documentation support, compilation is very fast, allowing a rapid write-compile-debug cycle. This speed of compilation gives rapid feedback on experimental approaches, improves language learning, and allows Go to be used in areas where scripting languages have been used because of their rapid development cycle.

Parallel computation has been noted as an important aspect of high performance computing in the context of genomic datasets (Knight *et al.*, 2007). The Go language has built-in support for concurrent processing, with primitives for initiating concurrent routines and allowing safe communication between them. This support eases implementation of parallel computation on appropriate architectures.

A significant difference between Go and other languages commonly used in bioinformatic computation, such as Python, Perl and Ruby is that Go is strongly typed. This strictness prevents a large class of programming errors by requiring explicit statement of programmer intention and by catching errors in code manipulation during the maintenance phase of program development. We believe that the explicit nature of strong typing also favours ease of peer review of published code. Subtle errors that do not cause program failure can be a significant cause for introduction of invalid data into the body of scientific literature (Wilson *et al.*, 2014); Go programming practice promotes the transparency in failure, failing ‘noisily’ and informatively rather than silently as is often the case with dynamically typed languages or uninformatively as is the case with C/C++. Go’s use of structural typing through interface types in conjunction with type composition via embedding retains the level of flexibility given by dynamic typing, and support for type inference reduces the burden on programmers to explicitly state types in all cases.

The Go standard library includes support for CPU and memory profiling to identify ‘hot’ or memory-expensive code that would benefit from optimization to improve performance. Further, the Go tool chain supports the inclusion of C and assembly code into packages, allowing an approach, similar to that used in PyCogent, when Go performance is inadequate.

In addition to this direct interaction between Go and C code, bíogo interfaces easily with external tools using the `biogo/external` package. This allows a concise description of command line parameters for an external application within the context of the Go type system. Defining new external application interfaces is a trivial exercise.

The bíogo library includes packages supporting common bioinformatic use cases: reading and writing sequence data in FASTA and FASTQ formats, and genomic interval data in BAM, BED and GTF formats; and storing metadata, sequence and interval data in a variety of general purpose data structures in addition to the built-in Go types. In addition to handling common data formats, bíogo supports Entrez and remote BLAST queries through the `biogo/ncbi` packages easing public data acquisition. Analytical tools are provided to support sequence alignment, data clustering, and numerical and graph analysis. In addition to the provided analytical tools, biogo provides support for graphical rendering of genomic data with a collection of graphics packages, including a package for generation of Circos-like plots (Krzywinski *et al.*, 2009), allowing direct integration with bíogo data types.

A significant feature of the bíogo library is its use of Go’s structural typing. An example of this is the interface relationships between genomic intervals defined in the `feat` package and sequence types defined in the **seq** package hierarchy; sequence types satisfy the feature interface, thus allowing sequences to be used in all situations where features are permitted (Examples in Supplementary material 1). Further, Go’s embedding and composition capacity allows easy construction of rich data types based on the provided bíogo types, for example extension of a sequence type to satisfy the `biogo/store/step` element interface allows a space-efficient storage and representation of interrupted chromosomal sequence using run-length encoding. Examples of the use of these language features in the context of the biogo library are provided in the `biogo/examples` and `biogo/talks` repositories, but we provide here some specific examples that demonstrate its utility in developing tools for analyzing large data sets.

We have used components of the bíogo library to implement a pure Go version of the genome-scale pairwise local sequence alignment tool, PALS (Edgar and Myers, 2005), as an example of a computationally intensive problem. We find that the performance of the Go PALS implementation is comparable with the C implementation and is more robust; on a workstation with a 2.7GHz Intel i7 processor and 12GB of RAM, processing of the 70Mbp *Oikopleura dioica* genome (obtained from the Oikpleura Genome Browser http://www.genoscope.cns.fr/externe/GenomeBrowser/Oikopleura/") with default PALS options is completed in 1 minute 25 seconds with the C implementation and 2 minutes 20 seconds with the Go implementation. Parallel processing is not possible with the C implementation, but is completed in 1 minute 40 seconds with the Go implementation. While this demonstrates a comparative reduction in computational performance, the Go implementation is performing additional work in ensuring memory safety where the C implementation does not. We have used this tool for analyzing retro-transposable repeats in mammalian genomes in place of C implementation of PALS due to the safety provided Lim *et al.* (2014).

**Figure 1:**
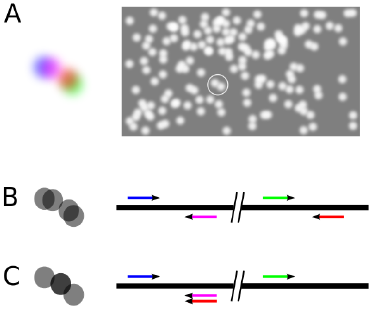
Possible impact of cluster collision on read pair mapping discordance. A. shows a diagram of a field of flow-cell image spots and a colliding pair of pair-end read pair spots (blue/purple and red/green); B. shows the true oracular identity of spots and resulting genomic mapping; and C. shows incorrect resolution, with discordant mapping due to one member of a pair (red) being attributed to the colliding pair mate (purple) image spot.

Another example is the analysis of Illumina HiSeq^TM^sequencing data to examine the effect of template cluster collision on paired-end read mapping discordance. We wanted to rule out cluster collision as a source of error in a structural variation analysis, so we designed an analysis to estimate the proportion of discordant read pair mappings due to incorrect image analysis by the Illumina sequencing pipeline (Figure 1). This involved finding the nearest neighbor spot of each discordantly mapped read pair and then finding its genomic mapping location, marking it as a possible cause if it had been mapped to the same genomic location as at least one mate of the discordant pair. This approach requires that the location and mapping data of all clusters of a flow-cell tile be stored to be able search for neighbors and compare mapping locations. We were able to implement this analysis in under 400 lines of source code (Supplementary material 2). An analysis of approximately 370 million paired-end 100nt read alignments from 91 flow-cell tiles using this code was completed in 12 hours with 90GB of system RAM allocating eight cores of a 2.3GHz AMD Opteron 6134. A smaller memory footprint would be possible if the mapping data were sorted by tile prior to the analysis such that only one tile need be kept in memory at a time. We found a statistically significant, but negligible effect of collision on read pair discordance (not shown).

Components of bíogo have been used in the implementation of a compressively accelerated protein BLAST which outperforms existing similar tools written in C (Daniels *et al.*, 2013). The CaBLASTP project makes use of Go’s built-in concurrency primitives to exploit multiprocessing CPUs.

## 3 Conclusion

bíogo is a simple, type-and thread-safe toolkit for the prototyping and development of bioinformatic tools. It is easy to learn and use, and aids in development of easily maintained and peer-reviewed code. Applications written using bíogo have comparable performance with tools written in C.

## 4 Supplementary Material

All bíogo packages and documentation are freely available from the web at https://github.com/biogo under a Modified BSD License.

## 5 Acknowledgments

We Would Like To Thank Russ Cox, Martin Frith and Dan Perkins for their feedback on the manuscript, and Stephen Bent for his input into the design of key aspects of the library. This work was supported by The University of Adelaide; and The National Health and Medical Research Council [grant number 1011334].

